# Algorithms in Stringomics (I): Pattern-Matching against “Stringomes”

**DOI:** 10.1101/001669

**Authors:** Paolo Ferragina, Bud Mishra

## Abstract

This paper reports an initial design of new data-structures that generalizes the idea of pattern-matching in stringology, from its traditional usage in an (unstructured) set of strings to the arena of a well-structured family of strings. In particular, the object of interest is a family of strings composed of blocks/classes of highly similar “stringlets,” and thus mimic a population of genomes made by concatenating haplotype-blocks, further constrained by haplotype-phasing. Such a family of strings, which we dub “stringomes,” is formalized in terms of a multi-partite directed acyclic graph with a source and a sink. The most interesting property of stringomes is probably the fact that they can be represented efficiently with compression up to their *k*-th order empirical entropy, while ensuring that the compression does not hinder the pattern-matching counting and reporting queries – either internal to a block or spanning two (or a few constant) adjacent blocks. The solutions proposed here have immediate applications to next-generation sequencing technologies, base-calling, expression profiling, variant-calling, population studies, onco-genomics, cyber security trace analysis and text retrieval.

## 1 Introduction and Motivations

One may define a “stringome” to be a family of strings that can be obtained by concatenation of a small number of shorter elemental strings – “stringlets,” which may additionally share many common structures, patterns and similarities or homologies. In particular, we may assume that these elemental strings can be partitioned into several classes such that the stringlets within a class are few in number and differ from each other by small edit-distances – thus giving rise to small number of “allelic forms” within each class. In a dynamic setting, the stringomes may evolve by adding, deleting or mutating the set of elemental strings, but also by changing the rules by which the strings may be concatenated in a stringome. Study of such combinatorial objects may then be referred to as “*stringomics*,” and raises many combinatorial and algorithmic questions related to efficient pattern matching, query processing and statistical analysis, which would otherwise be prohibitively expensive in the absence of any imposed structures and associated assumptions.

Thus, for instance, because of the constrained nature of these stringomes, one expects that they may be amenable to representation in a compressed data structure, while allowing searching, indexing and pattern-matching to be carried out in an efficient manner (possibly, directly in hardware). More importantly, in many domains, particularly in biology or linguistics, the stringomes can be used to directly represent and manipulate key sequence-information, which are known to be controlled by certain evolutionary dynamics. For instance, the framework proposed here provides a natural setting for many important computational problems related to genetic analysis of large number of genome sequences, which are typically presented with an imposed population stratifications and phenotypic annotations. Thus, the algorithmic studies in stringomics are directly relevant to such emerging fields as genomics, epigenomics, transcriptomics and proteomics – all capable of generating increasingly massive amount of raw data that promise to make a significant impact on human healthcare, agriculture, and manufacturing, e.g., green energy, pharmaceuticals, bio-remediable materials, etc. We also believe that suitable modifications to our approaches will eventually find applications to hypertext processing, natural language processing, speech processing, cyber security trace analysis, as well as other applications of similar nature.

We formulate the problems discussed below, motivated largely by our desire to immediately improve algorithmic, storage and data-transmission demands created by the state-of-the art genomics analysis, sequencing- and mapping-related biotechnologies, whole genome sequence assembly (WGSA) and genome-wide association studies (GWAS). Currently deployed genomics analysis algorithms are simple reincarnations of generic stringology algorithms that were developed in the most general unconstrained setting, and have failed to take advantage of the genome structure, e.g., what can be discerned in diploid eukaryotic genomes. For instance, while devising algorithms to study a population of human genomes, the computational genomicists have not yet taken noticeable advantage of the genome-architecture of such structural elements as haplotype blocks, sparsity in variations, allelic frequencies, haplotype phasing, population stratification, etc. New approaches that exploit these architectural structures to dramatically improve the algorithmic and communication complexity will undoubtedly benefit a plethora of genomic (as well as other-omics) applications: For instance, Bayesian base calling [24], genome assembly (both genotypic and haplotypic), resequencing, and most importantly, the embryonic field of clinical genomics (see [25]). Another specialized, but an enormously critical, application comes from the field of onco-genomics analysis, which studies the genomes of a heterogeneous population of tumor cells and stroma, all undergoing rapid somatic evolution, choreographed through cancer hallmarks, but also sculptured by the chemo- and radiation-based therapeutic interventions.

In summary, we believe that although the combinatorial objects, we study here—namely, stringomes—are somewhat specialized, they are likely to become foci of many future algorithmic innovations, since, just in the field of genomics, these will deliver many desperately needed tools to curb the data deluge, improve data-compression (both in transmission and storage), and ultimately, accelerate disease studies through intelligent pooling strategies (see e.g. [25]).

The paper is organized as follows. Section 2: historical notes and connection to pattern matching on hypertext; Section 3: a precise formulation of the Stringomes and the algorithmic pattern-matching problems related to them; Section 4 and 5: schematics of data structures constructed from String B-trees, Range-trees, FM-indices, Patricia tries, Geometric BWTs, etc. and complexity analysis for pattern-matching operations conducted using various combinations of them; and finally, Section 6: a more complex query system which restricts the stringomes to a suitable subset based on an embedding population-stratification structure.

## 2 Historical Notes

As it will be clarified in the next sections, our formalization of stringomics is reminiscent of certain concepts, developed to study “*pattern matching on hypertext*,” which was introduced by Manber and Wu in 1992 [23]. In that scenario, the hypertext was modeled in the form of a graph of *n* nodes and *m* edges, each node stores one string and edges indicate alternative texts-nodes that may follow the current text-node. The pattern is still a simple (linear) string of length *p*. A pattern occurrence was defined as a path of text-nodes which contain the pattern. Therefore, an occurrence may be thought of as *internal* to a text-node, or to span (a path of) several text-nodes.

Manber and Wu considered an acyclic graph and reported all *occ* pattern occurrences in *O*(*N + p m* + *occ* log log *p*) time, where *N* is the total length of the strings stored in the graph’s nodes. Akutsu [3] improved the solution for the case of a tree structure in optimal time, while Park and Kim [19] extended this result to an *O*(*N* + *pm*) time-algorithm for directed acyclic graphs (DAG) and for graphs with cycles where no text-node can match the pattern in two places. Subsequent to that, other researchers [1, 26, 27] dealt with the problem of *approximate* pattern matching on hypertexts, showing that the problem is NP-complete if the errors occur in the text, and they solved the problem in *O*(*pN*) time and *O*(*N*) space for cyclic and acyclic graphs.

Overall these are batch-solutions that need to scan the entire graph and the strings contained in its nodes in order to count/report the pattern occurrences. In this paper we aim at designing *index*-based solutions which can count/report the pattern occurrences in a time complexity which is as independent as possible to the graph and string sizes, while remaining succinct/compressed in the occupied space. Thus our solution must exploit the structural properties of stringomics problem which assumes that the stringlets within a class are few in number, and may differ from each other by small edit-distances and, in addition, the pattern queries may span only a few nodes (i.e., their number bounded by a constant, say, no more than 2). In future, we will extend the techniques to a much broader context, where no such bounds need be imposed.

## 3 Problem Statement

Our problem boils down to a special case of pattern matching on hypertext, in many respects. To make this connection explicit, we start with few general remarks: first the graph is directed and acyclic (DAG, Directed Acyclic Graph); second, the nodes can be partitioned into groups whose strings are highly similar to each other and thus can be highly compressed; third, pattern occurrences can span only a *constant* number of nodes, and for the sake of a simplified presentation, we assume hereafter that they can be at most two. This framework implies that a pattern occurrence can be either fully contained within a node-string, or it can overlap two node-strings connected by a “link” and represented by an edge, say (*s*′, *s*″), thus actually matching a suffix of *s*′ and a prefix of *s*″. In the rest of the paper, given the structure of these occurrences, we will call the former *node occurrences* and the latter *edge occurrences*. The case in which more than two nodes are involved in a pattern occurrence will be dealt in a future publication.

To be precise, our problem consists of *k* groups of variable-length strings `K`_1_, `K`_2_, …, `K`_*k*_, providing the building blocks for the “stringomes.” The strings are *n* in number, have a total length of *N* characters, and are further linked in a pair-wise fashion by *m* links, defined below more precisely. Each group `K`_*i*_ consists of *n*_*i*_ strings 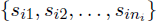, highly similar to each other. In many situations of practical interest to us, it could be assumed that |*s*_*ij*_| ≤ *S*_max_ and *n*_*i*_ is bounded from above by a small constant (e.g., in the genomics context, *S*_max_ ≈ 30*Kb*, the size of a typical human haplotype block and *n*_*i*_ ≤ 10, the number of different allelic forms for a haplotype block, though in principle there could be as many as 2^max|*s*_*ij*_|^ alleles, only accounting for biallelic single nucleotide polymorphisms or SNP’s). The indicator function, 𝟙_*s*′, *s*″_ is 1, if there is a link (edge) between the pair of strings (*s*′, *s*″) and 0, otherwise. It is, then, 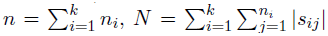, and 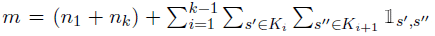. When the number of allelic forms are bounded by a small-constant, *n* = *O*(*k*), *m* = *O*(*k*) and *N* = *O*(*kS*_*max*_), thus making the complexity strongly dependent on the number *k* of haplotype blocks, and the size of the haplotype blocks *S*_max_, i.e., genome length and linkage dis-equilibrium (LD-) parameter. In species, such as *H.sapiens*, which is assumed to have encountered a recent population bottle-neck, these parameters are clearly advantageous to our algorithmic framework, as k is small and *s*_*ij*_’s are relatively long. Also, note that, given the strong group-similarity, the value of *N* is a pessimistic upper bound to the storage of the strings. We will deal with this issue next, and for the sake of presentation, the following bounds will be given in terms of the parameters *N* and *m*, resorting subsequently to the *k*-th order empirical entropy *H*_*k*_(`K`) of the string set `K` = U_*i*_ `K`_*i*_ when dealing with compressed data structures [30].

These groups of strings are interconnected to form a multi-partite DAG *G* = (*V*, *E*) defined as follows. The set *V* consists of *n* + 2 nodes, one node per string *s*_*ij*_ plus two special nodes, say s_0_ and *s*_*n*+1_, which constitute the “source” and the “sink” of the multi-partite DAG and contain empty strings (in order to avoid generating spurious hits). The set *E* consists of *m* edges which link strings of adjacent groups, namely we can have edges of the form (*s*_*ij*′_, *s*_(*i*+1)*j*″_), where 1 ≤ *j*′ ≤ *n*_*i*_ and 1 ≤ *j*″ ≤ *n*_*i*+1_. In addition, the source *s*_*0*_ is connected to all strings of group `K`_1_ and the sink *s*_*n*+1_ is linked from all strings in *`K`_k_*.

Figure 1 shows a pictorial example of this graph. In red are indicated the characters which are different from the ones in the corresponding position in the first string of each class. Bold green is used to color the occurrences of the pattern *P* = `accc`, indicating that we have 1 node-occurrence and 3 edge-occurrences, and these last type involves different splitting of the pattern between the two node-strings involved in the matching. We remark that our solutions work even in the case that edges connect strings of non-adjacent classes, namely *E* ⊆ *`K`_i_* × *`K`_j_* with *i < j*. However, in what follows, for the sake of simplicity of description, we will deal only with the class-adjacency assumption.

**Figure 1:**
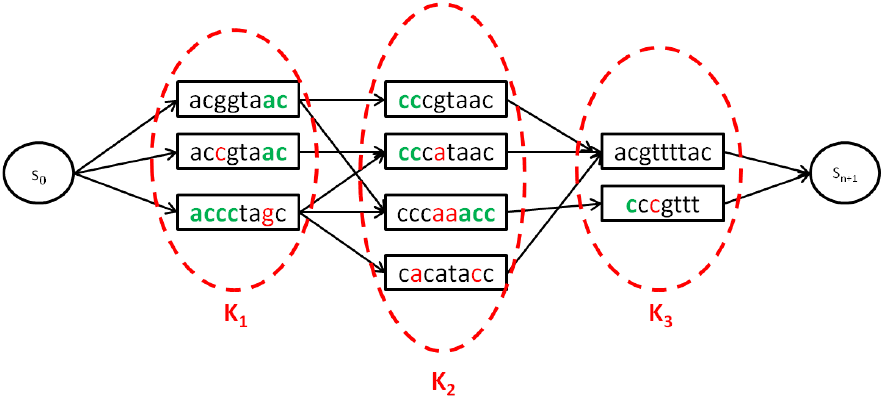
An example of DAG *G*. Note the occurrences of the pattern `accc`, highlighted in bold-green.

The main algorithmic question, we address in this paper, is the following: how can one build an index over *G* in order to efficiently support two basic pattern queries:

### Counting

Given a pattern *P*[1, *p*], we wish to count the *number occ* of pattern occurrences in *G*.

### Reporting

Same as the previous query, but here we wish to *report* the positions of these *occ* occurrences.^1^

Here, the challenging issue is that *G* encodes a multitude of strings, one per possible edge (or *path* in the more general case) in the DAG, so we cannot store all of them explicitly and then build a classic indexing data structure (such as suffix tree or array [15], or an FM-index [13, 30]), because that would result in the space growing as ⊝(*Nm*) rather than ⊝(*N + m*) in the worst case: recall that if one assumes *N* ≈ 10^10^ is the number of the characters in the stringomes and *m* ≈ 10^7^ is the total number of links defining the architecture of the stringomes, which are roughly the numbers for a large sample from the human genome population (e.g. HapMap population), then this complexity would make such a solution infeasible. In order to circumvent this problem, we need to account for the string interconnections at query time, rather than at indexing time, still guaranteeing that this step can be performed efficiently in time and space.

## 4 Schematics of a Data Structure

In this section we will describe the schematics of a data structure that underlie our algorithmic solution to the problem stated earlier. To achieve this goal, we will proceed as follows: first, identify the data-structural building blocks needed to answer the pattern queries, specify their general algorithmic features, and finally, detail some implementations for these data structures, which will allow us to derive the time/space bounds for our solutions. In particular, we will detail three kinds of implementations: one addressing the I/O-issue only, one addressing the compression-issue only, and the final issue combining the two. These implementations are best described as structured over a library of three kinds of data structures:

- `D`_K_ is a *full-text index*, efficiently supporting – both in time and space – substring searches over an indexed text, which can be built from the node strings suitably. `D`_K_ aims to find pattern occurrences residing totally within the graph nodes;
- `T`_K_ is a *dictionary index*, efficiently supporting – both in time and space – prefix searches over a dictionary of strings derived from the node strings and finally,
- `P`_K_ is a *geometric data structure*, efficiently supporting – both in time and space – 2d or 3d range queries over a set of points; these points are derived from the graph structure. In combination with `T`_K_, `P`_K_ facilitates finding the pattern occurrences crossing a limited number of edges/nodes in the graph.

This schematic approach is motivated by two reasons: (1) any advancements in the algorithmic community on any one of these data-structural problems can be deployed *as is* to improve correspondingly the results stated in this paper; (2) the expository structure of the paper has been simplified by distinguishing between the algorithmic solutions to our problems and their fine technical details, lest it would hinder its reading and understanding.

### 4.1 Node Occurrences

We create the string 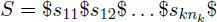 by concatenating all strings in `K` separated by a special character $ which is assumed to not occur in the strings’ alphabet. It follows that |*S*| = *O*(*N*).

Clearly if *P* is a node-occurrence in *G* then *P* occurs inside some string of `K`, and thus, by definition of string *S*, the pattern *P* occurs in this string. The converse is also immediately true. Therefore, counting/reporting node-occurrences of *P* in *G* boils down to counting/reporting pattern occurrences in *S*. Consequently we build a full-text index D_K_ over the string *S* which we assume to take *s*_*node*_(*S*) space and *tc*_*node*_(*S*, *p*) time to count the pattern occurrences of *P* in *S*, and *tr*_*node*_(*S*, *p*) time to report the pattern occurrences in *S*.

**Implementation.** `D`_K_ can be any known full-text index, such as the suffix tree or the suffix array [15]. In this case, it is *s*_*node*_(*S*) = *O*(*N* log *N*) bits of space, *tc*_*node*_(*S*, *p*) = *O*(*p* + log *N*) time to count the node occurrences, and *tr*_*node*_(*S*, *p*) = *tc*_*node*_(*S*, *p*) + *O*(*occ*) time to report the *occ* node occurrences.

Clearly, if *N* becomes larger than the internal memory *M*, then we need to resort to either an external-memory index or a compressed self-index. In the former case, we could use the String B-tree data structure [12] and thus take *tc*_*node*_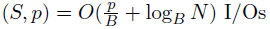 and *tr*_*node*_(*S*, *p*) = *tc*_*node*_(*S*, *p*) + 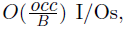, where B is the page size of the disk-memory system used to store the index and the string *S*.

In case a compressed self-index is used, such as the FM-index [13, 11] or the LZ-index [28, 20] or any of their variations [30, 29], then the counting time is *O*(*p*) and the reporting time per occurrence is slowed-down by a multiplicative term polylog(*N*) = *O*(log^i+ϵ^*N*/ log log *N*) or less depending on the implementation. The current literature is rich in variations of those compressed self-indexes, which offer several trade-offs between counting/reporting time and bits of compressed space, so we prefer to omit a detailed discussion of those variations, lest it would obfuscate the exposition. The reader could choose any of the known approaches (see e.g. [9, 30, 29]), as it would suit her performance needs, and incorporate them into our scheme to achieve suitable complexity bounds. Note that our solution implies (near) optimality in that, as far as the space is concerned, it achieves close to the k-th order entropy of the indexed string collection `K`, say *NH*_*k*_ (`K`)+ *o*(*N*) bits.^2^

An attentive reader might object that the set `K` is highly homogeneous in each sub-group `K`_*i*_ and yet the notion of entropy fails to capture this subtlety, since *H*_*k*_(*s*) ≈ *H*_*k*_(*ss*) and thus |*ss*|*H*_*k*_(*ss*) = 2|*s*|*H*_*k*_(*s*). As a result one might then mistakenly conclude that compressed indexes fail to take full advantage of the structure of `K`. However, it must be observed that the afore-mentioned space bounds are indeed upper bounds for the FM-index and the LZ-index. In practice those indexes achieve almost the same space for *s* and *ss*, as one can easily verify by running `bzip` or `gzip`. In any case, if one is interested in (also theoretically) better upper bounds in the space usage, then she may use the newly discovered family of self-indexes that are well-suited to the setting involving a collection of repetitive sequence and described in [22]. Their space occupancy depends only on the length of the underlying sequence (i.e. one of the strings in each group) and the number of variations in its repeated copies. This way the space reduction factor is no longer constant (and depending on the entropy of the underlying collection), but depends on *N*/*n*.

Recent progress in full-text indexing has achieved simultaneous improvements in both I/O-efficiency and succinct-space occupancy. State-of-the-art results use only *O*(*N*) bits of storage (here *σ* = *O*(1)), while supporting pattern searches either in *O*(*p*/(*B* log *N*) + (log^4^ *N*)/log log *n* + *occ* log_*B*_ *N*) I/Os [6] or in 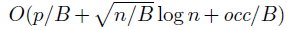 I/Os[16]. These improved complexities are due to a combination of geometric ideas and compressed full-text principles: called Geometric BWT [17].

### 4.2 Edge Occurrences, 1^st^ Solution

The non-triviality in our solution is in its ability to efficiently count/report the occurrences of *P*, which overlap the two strings at the extremes of a graph edge, say (*s*′, *s*″). Observe that such a pattern occurrence overlaps a suffix of *s*′ and a prefix of *s*″.

One solution consists of reducing this problem to a substring search problem in order to resort to a full-text index, again. We build a string *S*_*E*_ which is obtained by concatenating *m* substrings, which are obtained turning each edge *e* = (*s*′, *s*″) into the substring *s*_*e*_ = *s*′$*s*″$$. String *S*_*E*_ has length *O*(*mN*) in the worst case. This way, every pattern search boils down to executing *P* − 1 searches for patterns 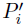 = *P* [1, *i*]$*P* [*i* + 1, *p*] for *i* = 1, 2, …, *p* − 1. It is clear that the counted/reported occurrences are only the edge-occurrences because the presence of the special symbol $ prevents returning node-occurrences which are internal in some string of `K`; whereas the presence of a double $$ between two strings generated by edges in *G* ensures that an edge-occurrence may occur only inside some *s*_*e*_. The advantage of this solution is in its ability to deploy known results in the (compressed) full-text index setting; the disadvantages are in the complexity: (*i*) the large space occupancy, exacerbated by the replication of each string as many times as the number of incident edges in E, and (*ii*) the slowdown in the query performance by a multiplicative factor *p*, induced by the searches for the 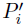s. On the other hand, in practice, the replication of the `K`’s strings may not be a problem if proper compressed data structures are used.

This approach thus takes *s*_*node*_(*S*_*E*_) space and *O(p × tc*_*node*_(*S*_*E*, *p*_)) time to count the edge occurrences in *G*, and *O(p × tr*_*node*_(*S*_*E*, *p*_)) time to report these edge occurrences.

**Implementation.** Given the reduction to full-text indexing, one may now resort to the ideas and solutions previously discussed in Section 4.1 and observe that the string *S*_*E*_ is even more repetitive than *S* but possibly much longer. So the space occupancy is close to |*S*_*E*_|*H*_*k*_(*S*_*E*_) bits, if a compressed self-index is used, whereas the time efficiency for counting the edge occurrences is *O*(*p*^2^) time, with an extra additive term of *O* (*occ* × polylog(*Nm*)) = *O* (*occ* × polylog(*N*)) time to report these occurrences. This approach has at least two appealing features, namely, its extreme simplicity in the way the compressed indexes are reused to search for both node- and edge-occurrences, which depends on how best to re-use known software libraries [11]; ease with which it can be parallelized given that the *p* searches can be executed independently of one another. On the other hand, its space utilization cannot be easily controlled and might grow explosively because of the necessity to explicitly create *S*_*E*_ before building the compressed index. Eventually external-memory algorithms to build the compressed full-text indexes could be used [10].

### 4.3 Edge Occurrences, 2^nd^ Solution

In order to avoid the replication of `K`’s strings we proceed differently and aim at using small space per edge, namely, space independent of the strings’ length. This solution reduces the edge-occurrence problem into a range-query problem over a set of *m* 2d-points, each one being in bijective correspondence with an edge of *G*.

We start by introducing two (sorted) sets of strings: `F` is the set of strings in `K` sorted alphabetically, `B` is the set of the reversal of the strings in `K`, also sorted alphabetically. We assign to each string *s*_*ij*_ ϵ `K` two ranks: the forward rank *fr*[*ij*] which indicates its (lexicographic) position in `F`, and the backward rank *br*[*ij*] which indicates the (lexicographic) position of its reversal in `B`. The simple key property that we will exploit is that, given a sorted set of strings, the ones prefixed by a given string occupy a contiguous range of positions. We can instantiate this property on the two sets `F` and `B` as follows: given the pattern *P*[1, *p*] and an integer i,

- the set of strings in `F` prefixed by *P*[*i* + 1, *p*] occupy a contiguous range of forward-ranks, say [*fw*_*start*_(*P* [*i* + 1, *p*]), *fw*_*end*_(*P* [*i* + 1, *p*])];
- the set of strings in `B` prefixed by (*P*[1, *i*])^*R*^ occupy a contiguous range of backward-ranks, say [*bw*_*start*_(*P*[1, *i*]), *bw*_*end*_(*P*[1, *i*])].

Now we notice that an edge-occurrence of *P* at the edge (*s*_*ij*_, *s*_(*i*+1)*j*′_) can be decomposed as a double match: one involves a pattern prefix *P*[1, *x*] and a suffix of *s*_*ij*_, and the other involves the remaining pattern suffix *P*[*x* + 1, *p*] and a prefix of *s*_(*i*+1)*j*′_. So we can partition the edge occurrences into *p* classes, say `P`_1_ ∪ … ∪ `P`_*p*_, where the class `P`_*x*_ contains all edges for which the matching pattern-prefix has length *x*.

Let us consider one of the edges in `P`_*x*_, say (*s*′, *s*″), then by definition *s*′ has suffix *P*[1, *x*] and *s*″ has prefix *P*[*x* + 1, *p*]; consequently, *s*′ has backward-rank in the sorted set `B` within the range [*bw*_*start*_(*P*[1, *x*]), *bw*_*end*_(*P*[1, *x*])], and *s*″ has forward-rank in the sorted set `F` within the range [*fw*_*start*_(*P*[*x*+1, *p*]), *fw*_*end*_(*P* [*x* + 1, *p*])].

As a result, we can retrieve all edge-occurrences `P`_*x*_ relative to the “string-pair” 〈*P*[1, *x*], *P*[*x*+1, *p*]〉 by just searching for all edges in *G* that satisfy both properties above. In order to solve this problem efficiently, we introduce an algorithmic transformation that maps the set *E* of *m* edges of *G* into a set `P` of *m* 2d-points as follows: each edge (*s*_*ij*_, *s*_(*i*+1)*j*′_) maps into the 2d-point (*bw*[*ij*], *fw*[(*i*+1)*j*′]).

Given this transformation, solving an edge-occurrence query boils down to a two-steps process:

**Prefix search:** For each prefix-suffix pair in *P*, namely 〈*P*[1, *x*], *P*[*x* + 1, *p*]〉, search for the range of strings in `F` prefixed by *P*[*x* + 1, *p*] and the range of (reversed) strings in `B` prefixed by (*P*[1, *x*])^*R*^. Let us denote the two string ranges by the ranges of forward/backward ranks delimited by that pair of strings: precisely, [*fw*_*start*_(*P*[*x*+1, *p*]), *fw*_*end*_(*P*[*x*+1, *p*])] and [*bw*_*start*_(*P*[1, *x*]), *bw*_*end*_(*P*[1, *x*])].
**Geometric search:** Solve a 2d-range query over the set `P` defined by [*bw*_*start*_(*P*[1, *x*]), *bw*_*end*_(*P*[1, *x*])] × [*fw*_*start*_(*P*[*x* + 1, *p*]), *fw*_*end*_(*P*[*x*+1, *p*])]. The counted/reported 2d-points in `P` are just the ones corresponding to the edges in *G* that identify an edge-occurrence, according to the way the set `P` is built and the geometric query is defined.

Overall the time cost to answer an edge-occurrence query boils down to *p* times the cost of a prefix-search over a set of *m* strings plus a range-query over a set of *m* 2d-points. It is clear that this approach has broken the dependency on *mN* which, instead, has characterized the previous solution in Section 4.2.

**Implementation of the prefix-search step.** The easiest and fastest way to solve the prefix-search problem over `F` and `B` is to build two compacted tries, one per set, taking *O*(*n* log *N* + *N*) bits of storage (recall that |`F`|= |`B`|= *n*). The time to prefix-search for a string *Q*[1, *q*] is *O*(*q*). This solution is efficient if space is not an issue and all computation fits in internal memory, otherwise the I/O-cost is ⊝(*q*) which becomes unacceptable given that the prefix-search must be repeated *p* times thus inducing eventually ⊝(*p*^2^) I/Os.

If I/O is a primary concern, then one could re-use the String B-tree data structure [12] storing the string set `F` and `B` (see Section 4.1) and take 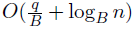 I/Os to identify the range of prefix-suffix occurrences.

Even in the most optimistic scenario, where all computation may fit in the internal memory, the space could remain a primary concern. If so, we could re-use the compressed self-index storing `K`, and supporting node-occurrence searches (see Section 4.1), and take *t*_*retrieve*_(*q*) = *O*(*q* + polylog(*N*)) time to retrieve any prefix/suffix of some string of `K`. Then we can build the Patricia trie data structure and adopt the *blind search* procedure, introduced in [12], to fast search for the prefix-suffix ranges in `F` and `B`. The choice of the Patricia trie, rather than the classic trie, offers two advantages:

- First, the space taken by Patricia tries is proportional to the number of indexed strings rather than their total length, given that they store just one integer per node and one character per edge. This way the space usage is *O*(*n* log *N*) bits rather than *O*(*n* log *N* + *N*) (indeed the indexed strings can be retrieved from the compressed self-index).
- Second, the prefix search can be implemented in *O*(*q* + *t*_*retrieve*_(*q*)) time and needs to access one single strings of `F` or `B`, rather than the previous *O*(*q*) accesses to strings (because of the *O*(*q*) edge traversals), which actually translate to ⊝(*q*) cache misses.

We denote these two Patricia tries by `T`_K_. They occupy *O*(*n* log *N*) bits of space and support a prefix/suffix search for a pattern *Q*[1, *q*] in *O*(*q* + *t*_*retrieve*_(*q*)) = *O*(*q* + polylog(*N*)) time.

If I/Os and space are a concern, then we could adopt a mixed solution that deploys the String B-tree to index a sample of `F` and `B`, and a classic compressor to store the strings in these two sets in a blocked fashion. The idea is to scan each set (either `F` or `B`) in lexicographic order and compress the strings one after the other until the compressed block has size ⊝(`B`).^3^ This process will continue to form at most *O*(*N*/*B*) blocks, possibly much less because of the compressibility of the sets `K`_*i*_ and the sorted order in which `F` and `B` are scanned. Actually one could argue that these blocks are *O*(*n*/*B*) because of the *O*(1) length of the indexed/compressed strings. Given this compressed blocking, the first string of each block is indexed in a String B-tree thus taking *O*(*n*/*B*) overall space. Searching for the prefix/suffix range in Fand Btakes *O*(*q*/*B* + log_*B*_(*n*/*B*)) = *O*(*q*/*B* + log_*B*_ *n*) I/Os.

Thus, as we had done for all previous data structures, we use the convention to denote by `T`_K_ the data structure used to index `F` and `B`, and by *t*_*range*_(*q*, `K`), the time complexity of determining the prefix/suffix ranges in `F` and `B`. Note that recently developed compressed string B-Tree [14] structure provides an ideal implementation of `T`_K_, which will be explored further in our implementation.

In [14], the prefix-search problem addressed by `T`_K_ has been solved optimally in space occupancy and I/Os, but more importantly, by relying only on a very general cache-oblivious model. These performance metrics are achieved, while matching the space lower bound for tries up to a constant multiplicative factor (1+ϵ), for ϵ > 0, and query complexity *O*(log_*B*_ *n* + (1+1/ϵ)*P*/*B*) I/Os. To avoid complexity, we only parenthetically mention this solution for `T`_K_, as, for instance, the theorem that follows below describes only a simpler solution (both easier to understand and implement) based on the plain String B-tree.

**Implementation of the geometric-search step.** To answer this query it is enough to build a (succinct) data structure for 2d-range queries over the set of points `P`. One implementation for this data structure is the classic Range Tree^4^ which takes *O*(*m* log *m*) space and *O*(log *m*) query time, since |`P`| = *m*. Counting time can be improved to 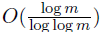 and space to *O*(*m*) [18]. Reporting can be speeded-up to *O*(log log *m*) time but taking *O*(*m* log^ϵ^ *m*) space (for any ϵ > 0), since the 2d-points have integer coordinates within a bounded space [4]. Space could be further reduced [7], by taking into account the statistical distribution of the points.

In case I/Os are a primary concern, we can employ an external-memory version of the Range-Tree [5, 2] and thus use *O*((*m*/*B*)(log *m*/ log log_*B*_ *m*)) disk pages and answer range queries with *O*(log_*B*_ *m* + *occ*/*B*) I/Os (in the case of counting, set *occ* = 0). There also exists a number of linear space solutions for 2d-range queries based on *k*d-trees, but with worse I/O-complexity.

we point out that compression here is not a primary issue because we do not know *a priori* the statistical distribution of the points (strings) and we assume that *m* ≪ *N*. In any case, we have abstracted away our strategy sufficiently to make it agnostic to the best available data structure for 2d-range reporting/counting, thus making it mainly a black-box tool for our solution. Thus, in the future, no matter whatever improvement will be made available in the literature, it can be economically exploited by our approach to improve all our bounds. Going forward, just as we did for all previous data structures, we assume to denote by `P`_K_ the data structure used to answer the 2d-range queries, and write *t*_*count*_(`P`) to denote the time cost of counting the number of points falling into a 2d-range, and write *t*_*report*_(`P`) to denote the time cost of reporting the points falling into a 2d-range. Some figures for these parameters have been given above.

#### Theorem 1

*Listed below are three possible implementations of our ensemble of data structures, which address three different contexts of use. We note parenthetically that these are not necessarily the best possible combinations but only offer a good trade-off between simplicity and efficiency*.

**I/O-efficiency:** *We use the following implementation, the String B-tree for* `D`_K_ *and for* `T`_K_, *the external-memory Range-Tree for* `P`_K_. *This implementation uses O*(*N*/*B*+(*m*/*B*)(log *m*/ loglog_*B*_ *m*)) *disk pages, which we can safely assume to be O*(*N*/*B*), *hence O*(*N* log *N*) *bits of space*.
**Compressed space:** *We use the following implementation, the FM-index for* `D`_K_, *two Patricia tries for* `T`_K_, *the Range-Tree for* `P`_K_. *This implementation uses NH*_*K*_(`K`) + *o*(*N*) + *n* log *N* + *m* log^2^ *m bits of space*.
**I/O + compression:** *We use the following implementation, the Geometric BWT for* `D`_K_, *the String B-tree for* `T`_K_, *a blocked compression scheme for the strings in `K`, an external-memory Range-Tree for* `P`_K_. *This implementation uses O*(*N* + *m* log *m*) *bits of space*.

## 5 Answering the Pattern Queries

We are now ready to implement our original counting and reporting queries for a generic pattern *P*[1, *p*] over the string-graph *G*. Here we comment on counting because it is our most prominent query (see its use in estimating conditional probabilities in a prior, e.g. [24], but also [21] for a direct application to resequencing). We leave to the reader the more straightforward details and the task of deriving the solution for the reporting query.

The counting of the node-occurrences can be done by querying the data structure `D`_K_, in *t*_*count*_(*p*) time, which is the time cost of counting the pattern occurrences in the string set `K`, ignoring the existence of the DAG.

The counting of the edge-occurrences, in accordance with the “2^nd^ solution,” requires to first compute the prefix-suffix matches for all “string-pair” 〈*P*[1, *x*], *P*[*x* + 1, *p*]〉 by deploying the data structure `T`_K_ and then executing the 2d-range queries on the identified *P* ranges of forward/backward ranks, and summing their countings. This process takes *O*(*p* × (*t*_*range*_(*p*, *K*) + *t*_*count*_(`P`))) time. It goes without saying that we could improve – arguably by little – this time complexity by taking advantage of better self-indexes or the fact that the prefix/suffixes we are searching for are prefixes/suffixes of the same string *P*. In practice we could even contemplate parallelizing the *P* searches. Nevertheless, as stated above, we omit these technicalities in order to keep the exposition transparent.

### Theorem 2

*By deploying the three implementations sketched in Theorem 1, we guarantee the following performance bounds*:

**I/O-efficiency:** *We use the following implementation, the String B-tree for* `D`_K_ *and for* `T`_K_, *the external-memory Range-Tree for* `P`_K_. *This implementation uses O*(*p* (*p*/*B* + log_*B*_ *n* + log_*B*_ *m*) + log_*B*_ *N*) *I/Os, which we can safely assume to be O*(*p* log_*B*_ *m* + log_*B*_ *N*) *I/Os because reasonably P* < *B and n* < *m*.
**Compressed space:** *We use the following implementation, the FM-index for* `D`_K_, *two Patricia tries for* `T`_K_, *the Range-Tree for `P`_K_. This implementation uses O*(*p* (*p* + polylog(*N*))) *time*.
**I/O + compression:** *We use the following implementation, the Geometric BWT for* `D`_K_, *the String B-tree for* `T`_K_, *a blocked compression scheme for the strings in `K`, an external-memory Range-Tree for* `P`_K_. *This implementation uses O*(*p* (*p*/*B* + log_*B*_ *n* + log_*B*_ *m*) + polylog(*N*))) = *O*(*p* log_*B*_ *m* + polylog(*N*))) *I/Os, because reasonably P* < *B* *and n* < *m*.

## 6 A More Sophisticated Query

Let us assume that, with the graph *G* and the strings `K`, we are also given a taxonomy `C` of classes to which the strings in `K` may belong to. In reality (e.g., in genetic contexts), each string *s* may be further assumed to belong to at most one class (leaf) *cl*(*s*) of `C` and, assuming identity by descent (IBD), it may then belong to all ancestors of that leaf in `C`. `C` is represented by a tree of arbitrary fan-out (in practice, one expects a small fan-out, e.g., 2), and thus *O*(*n*) in size. More precisely we could state that the number of leaves is ≤ max_*i*_ *n*_*i*_.

For the sake of exposition, we introduce an additional simplifying assumption constraining edges of *G* and classes of `C`, namely that for each edge (*s*′, *s*″) ϵ *E* the strings *s*′ and *s*″ belong to the same class in `C`. This constraint simplifies our description and might be relaxed as described by the remark at the end of this section.

The query we wish to support over the pattern *P* is still a counting/reporting query but with an additional constraint, namely that the counting/reporting is restricted to the strings of `K` which descend from a given class *cl* ϵ `C`. We remark that the queried *cl* can be any node of `C`, even internal. In the light of this query, the previous additional constraint is easily explained, it ensures that either *cl* includes both strings of an edge-occurrence, or it does not, in which case the edge-occurrence is dropped from the retrieval. The case in which the two strings of an edge belong to two different classes could introduce the cumbersome situation in which one string descends from *cl* but not the other. So to be an edge-occurrence, *cl* should be an ancestor of the classes of *s*′ and *s*″. This requirement seems stringent, but, as detailed next, not unmanageable. For expository reasons, we continue with the simpler formulation for a moment – just one class per edge.

In order to answer this new sophisticated query we need to model the additional constraint given by *cl*, and we deal with it by adding one additional dimension onto our geometric mapping which therefore ranges over the 3d-space (ℕ^3^) rather than the 2d-space (ℕ^2^). The idea is to map every edge into a 3d-point (rather than a 2d-point), and to correspondingly map every pattern-class query into a 3d-range query (rather than a 2d-range query). We implement this 3d-mapping by performing a Depth-First visit of `C` during which, at each class *u* ∈ `C`, we attach the pre-post numbers *pre*(*u*), *post*(*u*) which are computed during that visit. As is well-known, these two numbers satisfy a useful property: any descendant of *u* in `C` is assigned a range which is included in [*pre*(*u*), *post*(*u*)], and conversely, any ancestor of *u* in `C` is assigned a range that includes [*pre*(*u*), *post*(*u*)].

We then map each edge (*s*_*ij*_′) *s*_(*i*+1)*j*′_ into a 3d-point (*bw*[*ij*], *fw*[(*i* + 1)*j*′], *pre*(*c*)), where *c* is the class of `C` to which belong both strings of that edge. We could have used interchangeably *post*(*c*) = *pre*(*c*) + 1, because *c* is a leaf of `C`. We notice that the set `P` still consists of *m* points, which now map into a 3d-space (ℕ^3^). On this set we build a data structure that can solve efficiently counting-reporting queries over 3d-ranges: this data structure is an extension of the ones used in the 2d-case with a slight slowdown in the query time and small increase in the space usage. Indeed, the cost of adding the 3d-coordinate is just a multiplicative factor *O*(log *m*) in the query time and space complexity in internal memory, which becomes *O*(log_*B*_ *m* + *occ*/*B*) I/Os and *O*((*m*/*B*) (log *m*/log log_*B*_ *m*)^3^) disk pages in external memory [5].^5^ Needless to say, there do exist other tradeoffs/solutions but we prefer here, just as in the previous sections, to avoid technicalities and merely sketch the algorithmic description of our approach.

We are left with showing how to turn a pattern-class query 〈*P*, *cl*〉 into a 3d-range query in order to count correctly the edge-occurrences (reporting follows easily). As we did in the previous section, we map *P*’s query into a 2d-range query, and then we add the third constraint on the class *cl* by including the range [*pre*(*cl*), *post*(*cl*)] as the range in the third dimension of the query. The points of `P` which belong to this 3d-range denote actually the edge-occurrences whose strings descend from class *cl* (recall the nesting property of the pre-post ranges assigned by the DFS-visit).

As far as the node-occurrences are concerned, a new solution must be devised, different from the one introduced in Section 4. The problem in that case is that the pattern occurrences are reported in some arbitrary order, thus making it impossible to quickly select the ones which belong to strings in *cl*. Our new solution consists of writing down the strings of `K` in the order of the leaves of `C`, to which they belong, and with some special end-markers separating them, say $. This way, for any class *cl*, the strings that belong to *cl* are contiguous in that ordering. Let us call this ordered string *S*, which has a size *N*, of course. Then we index *S* via the data structure introduced in [32] which solves the so called *restricted-positional text indexing* problem: namely, given a pattern *P* and a range of positions (*a*, *b*) report/count all occurrences of *P* in *S*[*a*, *b*]. This way, given a counting/reporting query on *P* and the class *cl* we have to issue a constrained-positional query on *P* and the range of positions which include the leftmost and the rightmost strings, constrained to descend from *cl*. To store this pair of positions it is enough to keep two integers per `C`’s node, thus requiring *O*(*n*) space – overall. The only problem here is that at present the restricted-positional indexes known in the literature are not yet compressed and thus take space *O*(*N* log^ϵ^ *N*) bits, and answer the queries in *O*(*p* + *occ*) time. In some sense this is a sort of quasi-compressed solution, because of the sub-linearity in the number of words *N*, which would otherwise occupy ⊝(*N* log *N*) bits.

### Theorem 3

*By deploying the FM-index for* `D`_K_, *two Patricia tries for* `T`_K_, *the 3d Range-Tree for* `P`_K_, *and restricted-positional text index over S, we get a solution which occupies N*(*H*_*k*_(`K`) + log^ϵ^ *N*) + *m* log^3^ *m bits of space and uses O*(*p* (*p* + *polylog*(*N*))) *time to answer the pattern-class restricted queries*.

For the sake of practicality we suggest to use another approach which scales well with larger values of *N* and thus works nicely also in external memory. It consists of turning all node-occurrences of *G* into edge-occurrences: just split every string *s* ∈ `K` into pieces such that their length is surely shorter than a queried pattern. All these pieces are linked up in *G*. The number of edges now becomes *m* + *N*/*p*_min_, where *p*_min_ is a lower-bound on the length of the queried patterns. This splitting ensures that no node-occurrence cases will occur, since patterns are longer than *p*_min_. As a result, our solution boils down to counting edge-occurrences only and thus we can apply the previous 3d-mapping which still deploys compressed indexes to store the strings of `K` and the two tries of `T`_K_. The most space is however taken by the geometric data structure which answers the 3d-range queries and now indexes ⊝(*m* + *N*/*p*_min_) points (edges). Hence the space is not only dependent on *H*_*k*_(`K`), for the part of the data structure `T`_K_, but also on that term which grows linearly with the length of the strings in `K`. However, in practice, we can expect^6^ *p*_min_ > 30 so that we could argue that 1/*p*_min_ < *H*_*k*_(`K`) thus making the geometric data structure not much large.

### Theorem 4

*Let us set d* = ⊝(*m* + *N*/*p*_min_) *and assume P* > *p*_min_. *By deploying the String B-tree for* `T`_K_ *and the 3d Range-Tree for* `P`_K_, *we get an external-memory solution which occupies O*(*N*/*B* + (*d*/*B*)(log *d*/log log_*B*_ *d*)^3^)) *disk pages and O*(*p* (*p*/*B* + log_*B*_ *n* + log_*B*_ *d*)) = *O*(*p* log_*B*_ *d*) *I/Os to answer the pattern-class restricted queries, given that reasonably p* < *B and n* < *d*.

We make two remarks: (1) using the data structures of [14], one could improve the space complexity (e.g., weaken the contribution from the additive term *N*/*B*), and (2) by adopting the solution in footnote 5, one could reduce the space to have the form *O*(*d*/*B* (log *d*/ log log_*B*_ *d*)^2^)) but we would increase the I/Os by a multiplicative factor *O*(log *m*/ log log_*B*_ *m*). Following the strategy carried out through-out this paper, the reader could choose any of the known approaches, depending on what suits her performance needs, and incorporate them into our scheme to achieve suitable time/space/IO complexity bounds.

## 7 Conclusions

This paper has initiated the study of a new class of pattern matching problems over “Stringomes,” and proposed several algorithmic solutions, which are instantiations of a basic data-structural scheme. The resulting algorithms are shown to be rather simple and yet efficient in space and time, so they are amenable to be implemented by using known geometric and string-matching libraries (such as LEDA and PizzaChili, just to name a few) or as an extension of the FEMTO software package [8]. The solutions proposed here have immediate applications to next-generation sequencing technologies, base-calling, variant-calling, expression analysis, population studies and onco-genomics.

In the future we plan to investigate a dynamic setting, in which the stringomes may evolve by adding, deleting or mutating the set of elemental strings, but also by changing the rules by which the strings may be concatenated in a stringome. Moreover we will plan to experiment some of the proposed solutions over data coming from several domains of molecular biology as described below:

**Genomics:** Currently a vast amount of data are being generated from the genomic studies of organisms from related species, organisms from a single population but with a focus on polymorphic locations (e.g., Human Hapmap Project, Exom data from 1000 Genomes Project), or cells from a heterogeneous tumor undergoing somatic evolution. In each of these cases, our stringomic data structure and the related pattern-matching operations will be helpful in

1. developing better priors for clinical genomic tools (such as TotalReCaller),
2. dilution- or map-based approaches to haplotype sequence assembly (e.g., locations in stringomes obtained by TotalReCaller, arranged as an optimal directed path in the DAG; optimality chosen based on a technology-sensitive score function),
3. performing Genome Wide Association Studies (GWAS), carried out on-the-fly on the data structure without any explicit data-modeling or variant-calling steps (e.g., focusing only on tag-SNPs), and
4. comparing two populations (either of organisms or cancer cells from two patients).
**Epigenomics:** Many similar questions also arise in the context of epigenetic modifications (e.g., in terms of methylation sites) and can help in coding the organ specific variations in epigenomics among the cells from different tissue types.
**Transcriptomics:** Requiring much more immediate and critical attention from stringomics is the application area of RNASeq, which aims to create a precise map of all the transcripts encoded in a complex genome, e.g., mammalian genomes, and additionally, augmented with the measurements of all their isoform-by-isoform abundances. While RNAseq technology has led to some interesting algorithmic developments, e.g., Tophat, Cufflinks, Scriptures, etc., much remains to be done: Especially in

1. improvement in accuracies of base-calling in cDNA (TotalRecaller [24] for transcriptomes, which is complicated by intron-exon boundaries and alternate splicing),
2. discovery of unannotated transcripts, alternate splicing isoforms, noncoding or anti-sense transcripts, allelic variations within gene families (i.e., orthologs and paralogs) and
3. efficient and compressed storage and transmission of single-cell and temporally-varying transcriptomes (e.g., from circulating tumor cells). A core RNASeq application, we plan to implement, would require several steps to run – possibly, in an individualized manner: (1) development of mapped exons (including information from annotated genes, but also discovering unannotated exons, using TotalRecaller with sliding-windows mapped to reference-genome(s) as Bayesian priors), (2) a naïve stringomic organization of the exons that allow all possible splicing junctions, and (3) a statistical-refinement of the stringomic data-structures, based on online learning schemes from splice-junction reads. The result is expected to be very precise maps of the transcriptome of a complex genome that can be used directly by TotalReCaller [24] for realtime and personalized clinical transcriptomic applications (see [25]).
**Other -omics:** One expects other similar applications in the context of micro-RNAs, microbiomics, and possibly, proteomics. Thus, the introduction of the concept of stringomic data-structures will fundamentally change how we have formulated the basic biological questions.
**General:** In general, the Stringomics data structures allow very flexible and non-parametric encodings of (empirical) Bayesian priors that could be generated from available data, and thus are likely to become critical to all Data Science applications, where data could be unstructured, but representable as strings, or their higher-order analogs.

### Note Added in Proof

It has been brought to our attention that, in a recent paper [31], very similar algorithmic ideas have been proposed to solve the main stringomics problem of §3. Nonetheless, we believe that our exposition (especially with the biological motivations) may still be relevant to our readers interested in various -omics applications, and that the implementation details we provide (in terms of I/Os and/or compression) may still be beneficial to the software engineers interested in creating a pipeline out of the state-of-the-art building blocks (e.g., `D`_K_, `T`_K_ and `P`_K_). In particular, discussions in §4.2 are simple but important for good practical implementation, and discussions in §6 deal with a new problem (motivated by population stratification studies) and an original solution.

## Acknowledgments

We wish to express our deep gratitude to Prof. Alfredo Ferro, who over the last several decades has remained a close intellectual friend of the NYU/Courant Bioinformatics Laboratory, has been a source of many stimulating research collaborations and played an instrumental role in catalyzing PF’s visit to the Courant institute. We also wish to thank those members of the NYU/Courant Bioinformatics Laboratory and MRTechnology, who contributed enormously in shaping this research: They are Will Casey, Mike Ferguson, Fabian Menges, Harsh Patel, Chang Peng, Jason Reed and Andi Witzel.

## Footnotes

1 It is clear that the identification of a pattern occurrence may involve in our DAG setting three integers: one to identify the source string, (optional) one to identify the destination string, and one to specify the offset of the pattern occurrence in the source string.

2 We state here that the space bound of the compressed indexes includes an extra additive term of *o*(|*S*| log *σ*) bits, where *σ* is the alphabet size, which is in the bio-setting a small constant, e.g., 4 (bases) or 20 (amino acids).

3 Here we could anyway guarantee that the `K`_*i*_’s strings are close to each other and fit in a block, or a contiguous sequence of blocks, so taking full advantage of the block compression.

4 http://en.wikipedia.org/wiki/Range_tree

5 Another interesting trade-off is *O*(*m*/*B* (log *m*/log log_*B*_ *m*)^2^) space and answers 3d-range queries in *O*(log_*B*_ *m*/(log *m*/log log_*B*_ *m*) + *occ*/*B*) I/Os.

6 For instance, using the currently popular Illumina MySeq sequencing platform, one may assume that the first 30-45 bases called will be subject to an error rate below 1%.

## References

[1] N. L. A. Amir, M. Lewenstein. Pattern matching in hypertext. In Proc. WADS, Lecture Notes in Computer Science, Vol. 1272, pages 160–173, 1997.

[2] P. Afshani, L. Arge, and K. Larsen. Orthogonal range reporting in three and higher dimensions. In IEEE FOCS, pages 149–158, 2009.

[3] T. Akutsu. A linear time pattern matching algorithm between a string and a tree. In Proc. CPM, Lecture Notes in Computer Science, Vol. 1272, pages 1–10, 1993.

[4] S. Alstrup, G. S. Brodal, and T. Rauhe. New data structures for orthogonal range searching. In Proc. FOCS, pages 198–207, 2000.

[5] L. Arge and K. G. Larsen. I/o-efficient spatial data structures for range queries. SIGSPATIAL Special, 4(2):2–7, 2012.

[6] Y.-F. Chien, W.-K. Hon, R. Shah, and J. S. Vitter. Geometric burrows-wheeler transform: Linking range searching and text indexing. In Procs of the Data Compression Conference (DCC), pages 252–261. IEEE Computer Society, 2008.

[7] A. Farzan, T. Gagie, and G. Navarro. Entropy-bounded representation of point grids. In Proc. ISAAC, volume 6507 of Lecture Notes in Computer Science, pages 327–338, 2010.

[8] M. P. Ferguson. Femto: Fast search of large sequence collections. In CPM, pages 208–219, 2012.

[9] P. Ferragina. *Handbook of Computational Molecular Biology*, chapter Chap. 35: String search in external memory: algorithms and data structures. Chapman & Hall/CRC Computer and Information Science Series, 2005.

[10] P. Ferragina, T. Gagie, and G. Manzini. Lightweight data indexing and compression in external memory. Algorithmica, 63(3):707–730, 2012.

[11] P. Ferragina, R. González, G. Navarro, and R. Venturini. Compressed text indexes: From theory to practice. ACM Journal of Experimental Algorithmics, 13, 2008.

[12] P. Ferragina and R. Grossi. The string B-tree: A new data structure for string search in external memory and its applications. Journal of the ACM, 46(2):236–280, 1999.

[13] P. Ferragina and G. Manzini. Indexing compressed text. Journal of the ACM, 52(4):552–581, 2005.

[14] P. Ferragina and R. Venturini. Compressed cache-oblivious string b-tree. In ESA, pages 469–480, 2013.

[15] D. Gusfield. Algorithms on Strings, Trees, and Sequences - Computer Science and Computational Biology. Cambridge University Press, 1997.

[16] W.-K. Hon, R. Shah, S. V. Thankachan, and J. S. Vitter. On entropy-compressed text indexing in external memory. In Procs of the SYmposium on String Processing and Information Retrieval (SPIRE), volume 5721 of Lecture Notes in Computer Science, pages 75–89. Springer, 2009.

[17] W.-K. Hon, R. Shah, and J. S. Vitter. Compression, indexing, and retrieval for massive string data. In Procs of Symposium on Combinatorial Pattern MAtching (CPM), volume 129 of Lecture Notes in Computer Science, pages 260–274. Springer, 2010.

[18] J. Jájá, C. W. Mortensen, and Q. Shi. Space-efficient and fast algorithms for multidimensional dominance reporting and counting. In Proc. ISAAC, volume 3341 of Lecture Notes in Computer Science, pages 558–568, 2004.

[19] D. K. K. Park. String matching in hypertext. In Proc. CPM, Lecture Notes in Computer Science, Vol. 937, pages 318–329, 1995.

[20] S. Kreft and G. Navarro. Self-indexing based on lz77. In Proc. CPM, volume 6661 of Lecture Notes in Computer Science, pages 41–54, 2011.

[21] B. Langmead. Highly Scalable Short Read Alignment with the Burrows-Wheeler Transform and Cloud Computing. M.S. Thesis, University of Maryland, College Park, 2009.

[22] V. Mäkinen, G. Navarro, J. Sirén, and N. Välimäki. Storage and retrieval of individual genomes. In Proc. RECOMB, volume 5541 of Lecture Notes in Computer Science, pages 121–137, 2009.

[23] U. Manber and S. Wu. Approximate string matching with arbitrary costs for text and hypertext. In Proc. IAPR Workshop on Structural and Syntactic Pattern Recognition, pages 22–33, 1992.

[24] F. Menges, G. Narzisi, and B. Mishra. Total recaller: improved accuracy and performance via integrated alignment and base-calling. Bioinformatics, 27(17):2330–2337, 2011.

[25] B. Mishra. The genome question: Moore vs. jevons. Jnl. of Computing of the Computer Society of India, 2012.

[26] G. Navarro. Improved approximate pattern matching on hypertext. In Proc. LATIN, Lecture Notes in Computer Science, Vol. 1380, pages 352–357, 1998.

[27] G. Navarro. Improved approximate pattern matching on hypertext. Theoretical Computer Science, 237(1-2):455–463, 2000.

[28] G. Navarro. Implementing the lz-index: Theory versus practice. ACM Journal of Experimental Algorithmics, 13, 2008.

[29] G. Navarro. Wavelet trees for all. In Proc. of the Symposium on Combinatorial Pattern Matching (CPM), volume 7354 of Lecture Notes in Computer Science, pages 2–26. Springer, 2012.

[30] G. Navarro and V. Mäkinen. Compressed full-text indexes. ACM Computing Surveys, 39(1), 2007.

[31] C. Thachuk. Indexing hypertext. J. Discrete Algorithms, 18:113–122, 2013.

[32] C.-C. Yu, B.-F. Wang, and C.-C. Kuo. Efficient indexes for the positional pattern matching problem and two related problems over small alphabets. In Proc. ISAAC, volume 6507 of Lecture Notes in Computer Science, pages 13–24, 2010.

